# ELAPOR1 induces the classical/progenitor subtype and contributes to reduced disease aggressiveness through metabolic reprogramming in pancreatic cancer

**DOI:** 10.1101/2023.09.23.558894

**Authors:** Yuuki Ohara, Amanda J. Craig, Huaitian Liu, Shouhui Yang, Paloma Moreno, Tiffany H. Dorsey, Helen Cawley, Azadeh Azizian, Jochen Gaedcke, Michael Ghadimi, Nader Hanna, Stefan Ambs, S. Perwez Hussain

## Abstract

Pancreatic ductal adenocarcinoma (PDAC) is a heterogeneous disease with distinct molecular subtypes classified as classical/progenitor and basal-like/squamous. We hypothesized that integrative transcriptomic and metabolomic approaches can identify candidate genes whose inactivation contributes to the development of the aggressive basal-like/squamous subtype. Using our integrated approach, we identified endosome-lysosome associated apoptosis and autophagy regulator 1 (ELAPOR1/KIAA1324) as a candidate tumor suppressor in both our NCI-UMD-German cohort and validation cohorts. We found that decreased ELAPOR1 expression was significantly associated with high pathological grade, advanced disease stage, the basal-like/squamous subtype, and decreased survival in PDAC patients. *In vitro* experiments showed that ELAPOR1 transgene expression inhibited migration and invasion of PDAC cells. Metabolomic analysis of patient tumors and PDAC cells revealed a metabolic program associated with both upregulated ELAPOR1 and the classical/progenitor subtype, encompassing upregulated lipogenesis and downregulated amino acid metabolism. 1-methylnicotinamide, an oncometabolite derived from S-adenosylmethionine, was inversely associated with ELAPOR1 expression and promoted migration and invasion of PDAC cells *in vitro*. Taken together, our data suggest that enhanced ELAPOR1 expression promotes transcriptomic and metabolomic characteristics that are indicative of the classical/progenitor subtype, whereas its reduction associates with basal-like/squamous tumors with increased disease aggressiveness in PDAC patients. This positions ELAPOR1 as a promising candidate for diagnostic and therapeutic targeting in PDAC.

**Novelty and Impact:** Pancreatic ductal adenocarcinoma (PDAC) exhibits heterogeneous molecular subtypes: classical/progenitor and basal-like/squamous. Comprehensive transcriptome and metabolome analyses in the PDAC patient cohorts and PDAC cell lines revealed that elevated ELAPOR1 correlates with enhanced survival, reduced PDAC cell invasion, and a distinct metabolic signature resembling the classical/progenitor subtype. Additionally, 1-methylnicotinamide has been identified as an oncometabolite, showing an inverse correlation with ELAPOR1. These findings emphasize ELAPOR1’s potential as a diagnostic and therapeutic target in PDAC.

**Graphical abstract:** 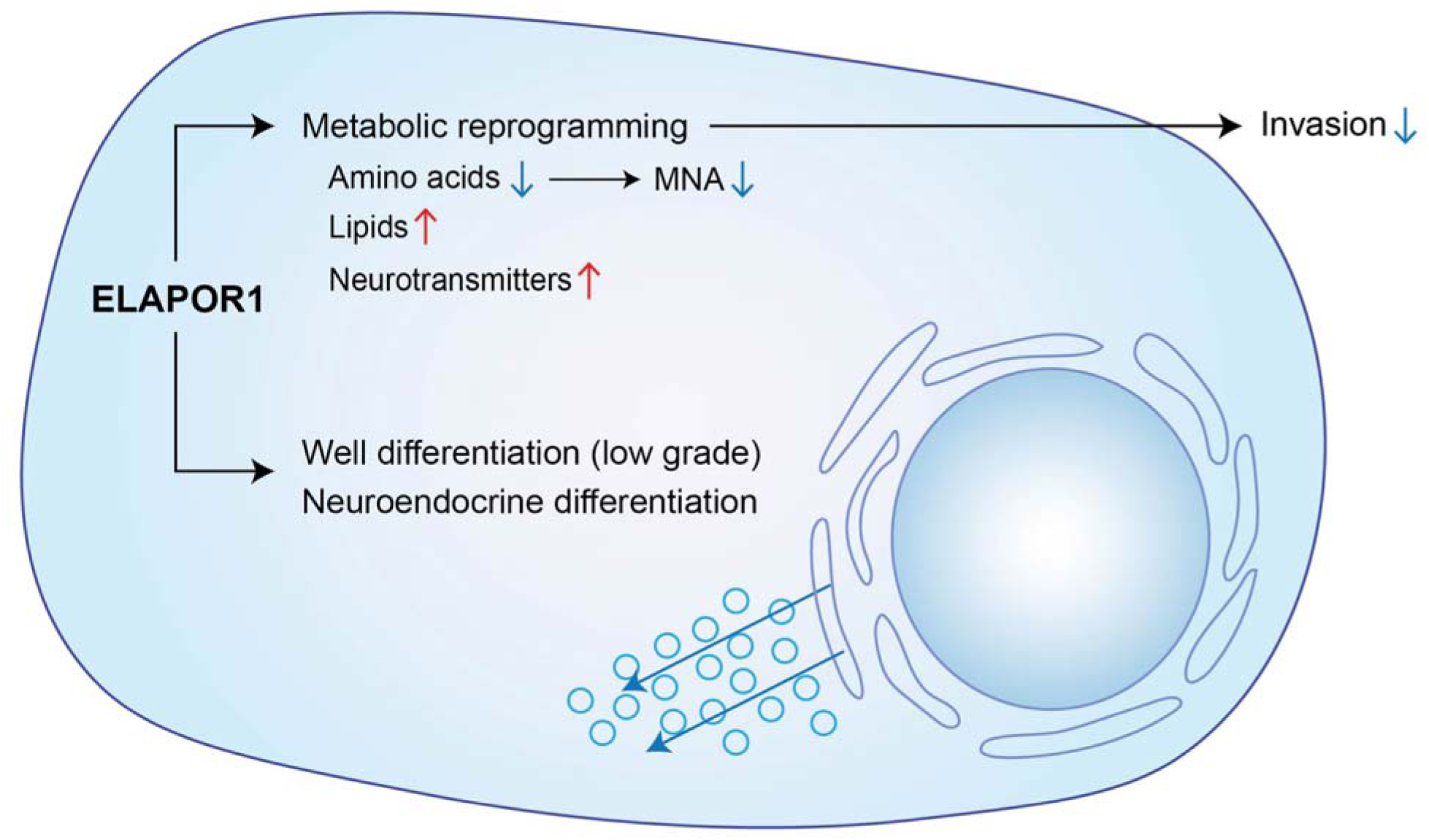

**Highlights:**

- ELAPOR1 is downregulated in basal-like/squamous PDAC
- Upregulation of ELAPOR1 associates with improved PDAC survival and reduced migration and invasion in PDAC cells
- ELAPOR1 expression induces a distinct metabolic signature as characterized by upregulation of lipogenesis and downregulation of amino acid metabolism, commonly observed in the classical/progenitor PDAC subtype
- The oncometabolite, 1-methylnicotinamide (MNA) is decreased when ELAPOR1 is upregulated, and promotes the migration and invasion of PDAC cells

## Introduction

Pancreatic cancer is a highly lethal cancer with a 5-year survival of only 12%^1^. Pancreatic ductal adenocarcinoma (PDAC) is the most common type of pancreatic cancer, accounting for more than 95% of all malignancies in the pancreas^2^. Earlier studies identified several molecular subtypes of PDAC with differences in biological characteristics and patient survival^3–5^. *Bailey* et al. proposed a classification system for PDAC based on four subtypes: pancreatic progenitor, aberrantly differentiated endocrine exocrine (ADEX), squamous, and immunogenic^3^. Both pancreatic progenitor and ADEX subtypes upregulate transcriptional networks associated with pancreatic development/differentiation of exocrine and neuroendocrine lineages^3^. The squamous subtype is associated with the most aggressive disease and exhibits distinct characteristics, including hypoxia, squamous differentiation, and MYC activation^3^. A more comprehensive subtype analysis recently proposed that PDAC can be classified into two major subtypes: ‘classical/progenitor’ and ‘basal-like/squamous’^6^. Beyond this distinction, the classical/progenitor subtype possesses more of an endodermal-pancreatic identity, while it is lost in the basal-like/squamous subtype^7^.

Metabolic reprogramming is one of the hallmarks of cancer^8–10^. PDAC rewires metabolism as part of tumor progression^10–12^. Moreover, the molecular subtypes of PDAC exhibit distinctively different metabolic profiles. The classical/progenitor subtype may be more dependent on lipogenesis for its metabolic needs whereas the basal-like/squamous subtype relies mostly on glycolysis^13–16^. However, the metabolic adaptations and their potential interaction with tumor gene-expression profiles have not been well studied in PDAC. Many of the previous studies were based on a small number of patient samples and cell lines. Using large PDAC patient cohorts together with human PDAC cell lines, we recently described SERPINB3 as an oncogenic driver of aggressive basal-like/squamous subtype of PDAC^17^. In the present study, we aimed to examine the interactive role of potential tumor suppressor genes and metabolic reprogramming in the development of molecular subtypes in PDAC. Our findings showed that downregulation of endosome-lysosome associated apoptosis and autophagy regulator 1 (ELAPOR1, also known as KIAA1324/EIG121) contributes to the metabolic adaptation in the basal-like/squamous subtype in patients with PDAC.

## Materials and Methods

### PDAC cohorts

We utilized gene expression datasets from the “Bailey” cohort (GSE36924)^3^, the “Moffitt” cohort (GSE71729)^4^, and our NCI-UMD-German cohort (GSE183795)^18^ to identify genes involved in the development of PDAC molecular subtypes. Partek Genomics Suite 7.0 (Partek Inc., Chesterfield, MO) was used to compare the basal-like/squamous subtype with the classical/progenitor subtype. For the integrative transcriptomic and metabolomic analyses, we used RNA sequencing data from patient PDAC tumors in the NCI-UMD-German cohort (GSE224564)^17^.

### RNA sequencing

RNA sequencing data were obtained by performing quadruplicate RNA sequencing using total RNA isolated from human PDAC cell line (Panc 10.05 ± ELAPOR1). The PDAC cells were cultured in RPMI 1640 medium, GlutaMax^TM^, supplemented with 10% FBS and 1% penicillin–streptomycin for 72 hours before RNA extraction. Libraries were prepared by the Sequencing Facility at NCI-Leidos using the TruSeq Stranded mRNA Kit (Illumina, San Diego, CA) and sequenced paired-end on NextSeq (Illumina) with 2 x 101 bp read lengths, as described earlier^17^. Briefly, a total of approximately 32 to 60 million paired-end reads were generated with a base call quality of ≥ Q30. Sequence reads in fastq format were then aligned to the human reference genome hg38 using STAR and RSEM to obtain gene expression as transcript per million with FPKM mapped reads. Differential expression analysis was performed using DESeq2. Ingenuity pathway analysis (IPA, QIAGEN, Venlo, Netherlands) and Gene Set Enrichment Analysis (GSEA) were used for the enrichment analysis in archived pathways and datasets. The RNAseq data for human PDAC cell lines were deposited in the NCBI’s GEO database under accession number GSE224417. The sequencing coverage and quality statistics for each sample are summarized in Table S1.

### Quantitative real-time PCR for ELAPOR1

The High-Capacity cDNA Reverse Transcription Kit (Thermo Fisher Scientific, Waltham, MA) was used to create first-strand cDNA from total RNA. Quantitative RT-PCR (qRT-PCR) assays were then performed using Taqman probes (Thermo Fisher Scientific): *ELAPOR1* (Hs00331399_m1) and *GAPDH* (Hs99999905_m1).

### Metabolic profiling and data analysis of PDAC

Metabolic profiling of the tumor samples was conducted by Metabolon Inc. (Morrisville, NC), using their standard protocol as described earlier^19–21^. Briefly, Metabolon’s untargeted metabolic platform uses two separate ultra-high performance liquid chromatography/tandem mass spectrometry (UHPLC/MS/MS) injections and one gas chromatography/mass spectrometry injection for each sample, to measure all metabolites. The resulting dataset was normalized across samples prior to data delivery, using a Metabolon protocol. The normalized relative abundance levels for each metabolite were used for further data analysis. The same protocol was used to perform metabolic profiling of cultured cells. Here, we performed quadruplicate metabolic profiling of human PDAC cell line (Panc 10.05 ± ELAPOR1). The PDAC cells were cultured in RPMI 1640 medium, GlutaMax^TM^, supplemented with 10% FBS and 1% penicillin– streptomycin for 72 hours. The pellets were collected, stored at −80 °C, and delivered to Metabolon Inc. for the UHPLC/MS/MS. IPA and MetaboAnalyst 5.0 (https://www.metaboanalyst.ca) were used for the enrichment analysis.

### Cell lines and culture condition

Human pancreatic cancer cell lines were purchased from American Type Culture Collection (ATCC), Rockville, Maryland. All cell lines were authenticated using short tandem repeat (STR) profiling within the last three years. All experiments were performed with mycoplasma-free cells. The CFPAC-1 (RRID:CVCL_1119) and Capan-1 (RRID:CVCL_0237) cell lines were cultured in IMDM medium supplemented with 10% FBS and 1% penicillin-streptomycin. The Capan-2 (RRID:CVCL_0026) cell line was maintained in McCoy’s 5A (Modified) medium, also supplemented with 10% FBS and 1% penicillin-streptomycin. The remaining PDAC cell lines (AsPC-1; RRID:CVCL_0152, BxPC-3; RRID:CVCL_0186, MIA PaCa-2; RRID:CVCL_0428, PANC-1; RRID:CVCL_0480, Panc 10.05; RRID:CVCL_1639, SU86.86; RRID:CVCL_3881) were cultured in RPMI 1640 medium with GlutaMaxTM, 10% FBS, and 1% penicillin-streptomycin, all in a humidified incubator with 5% CO2 at 37°C. All reagents for cell culture were purchased from Thermo Fisher Scientific.

### ELAPOR1 overexpression after lentiviral infection

The ELAPOR1 construct (EX-L2758-Lv122) and the corresponding empty vector control (EX-NEG-Lv122) were purchased from Genecopoeia (Rockville, MD). To establish stable cell lines overexpressing ELAPOR1, PDAC cells (Panc 10.05; SU86.86) were infected with lentiviral particles produced by transfecting 293T cells with the lentiviral expression vectors and the Lenti-Pac^TM^ HIV Expression Packaging system from Genecopoeia. Stable clones of the PDAC cells were obtained by selection with 4 μg/ml puromycin (Thermo Fisher Scientific).

### Cell proliferation assay

PDAC cells (Panc 10.05 cells, SU86.86 cells) were seeded in a 96-well plate to perform the CCK-8/WST-8 assay. The proliferation assay was conducted for 24, 48, 72, and 96 hours after seeding, following the manufacturer’s protocol (Dojindo Laboratories, Kumamoto, Japan). The absorbance was measured using a SpectraMax® ABS Plus microplate reader (Molecular Devices, San Jose, CA).

### Cell migration and invasion assay

Migration assay was performed with 24-well Falcon^®^ Cell Culture Insert (Corning, Glendale, AZ). For invasion assay, Matrigel Basement Membrane Matrix (#354234) was purchased from Corning. We coated the membrane of the upper chamber with 100 μl of Matrigel matrix coating solution (Matrigel matrix: coating buffer = 1:39) for 2 hours according to the manufacturer’s protocol. 750 μl of RPMI 1640, GlutaMax^TM^ supplemented with 10% FBS were added to the lower chamber and PDAC cells (Panc 10.05; 10×10^4^ cells, SU86.86 5×10^4^ cells) in 500 μl serum-free RPMI 1640, GlutaMax^TM^ were loaded into each upper chamber. The cells were incubated for 48 hours in a humidified incubator containing 5% CO_2_ at 37°C. After incubation, cells that had migrated or invaded through the membrane were fixed with 100% methanol (Thermo Fisher Scientific) and stained with Crystal violet solution (MilliporeSigma, Burlington, MA). The cells were then counted. To examine the effect of 1-methylnicotinamide (MilliporeSigma, Burlington, MA), 10 mM of the compound (final concentration) was added into the lower chamber.

### Immunohistochemistry

Four μm thick paraffin-embedded tumor sections and adjacent nontumor sections were incubated with rabbit monoclonal anti-ELAPOR1 antibody (MilliporeSigma, SAB2900912, 1:1000) overnight at 4°C. Signals were amplified using the Dako envision+ system-HRP labeled polymer anti-rabbit antibody (Agilent Technologies, Santa Clara, CA). Color development was performed with diaminobenzene (DAB, Agilent Technologies). Immunostaining was evaluated by assigning the intensity and prevalence score as described elsewhere^22, 23^. A score of 0–3, representing either negative, weak, moderate, or strong expression was assigned for intensity and prevalence was given a score of 0–4, representing <10%, 10–30%, >30–50%, >50–80% and >80% cells showing ELAPOR1 expression. Then we obtained the overall IHC score by multiplying the intensity and the prevalence score.

### Immunofluorescence

1.0×10^4^ of PDAC cells were seeded on 8-well Nunc^TM^ Lab-Tek^TM^ II CC2^TM^ Chamber Slides (Thermo Fisher Scientific). Cells were washed with PBS and fixed in 4% paraformaldehyde in PBS at room temperature for 10 min, rinsed twice and permeabilized with 0.2% Triton X-100 in PBS for 20 minutes. Cells were rinsed again with PBS and blocked in Animal-Free Blocker® and Diluent, R.T.U. (Vector Laboratories, Inc., Newark, CA; SP-5035-100) for 2 hours at room temperature before overnight incubation with rabbit polyclonal anti-ELAPOR1 antibody (Thermo Fisher Scientific, PA5-72691, 1:150). Cells were then washed three times with PBS for 15 min and incubated with anti-rabbit IgG (H+L) highly cross-adsorbed Alexa Fluor 594 secondary antibody (Thermo Fisher Scientific, A-21207, 1:2000) for 1 hr in the dark and at room temperature. Cells were then washed for 15 min, cultured with DAPI (Thermo Fisher Scientific, 62248, 1:1000) for 10 min, and mounted using ProLong™ Glass Antifade Mountant (Thermo Fisher Scientific). All samples were photographed at identical exposure times.

### Immunoblotting

To extract proteins, PDAC cells were lysed with RIPA Lysis and Extraction Buffer (Thermo Fisher Scientific). The protein extracts were electrophoresed under reducing conditions on 4– 15% polyacrylamide gels (Bio-Rad Laboratories, Inc., Hercules, CA) and transferred onto a nitrocellulose membrane (Bio-Rad Laboratories, Inc.). The membrane was incubated with SuperBlock™ Blocking Buffer (Thermo Fisher Scientific) for 1 hour at room temperature and then incubated with primary antibody overnight at 4°C. The following primary antibodies were used: ELAPOR1 (MilliporeSigma, SAB2900912, 1:500) and β-Actin (MilliporeSigma, A5441, 1:2000). The membrane was then incubated with secondary ECL anti-rabbit or anti-mouse IgG HRP-linked antibody (GE Healthcare, Pittsburgh, PA) for 1 hour at room temperature. Finally, we visualized the protein using a SuperSignal™ West Dura Extended Duration Substrate (Thermo Fisher Scientific).

### Statistical analysis

Statistical analyses were performed using GraphPad Prizm 9 (GraphPad Software, La Jolla, CA, USA). To determine the difference in overall survival between patient groups, we used the Kaplan-Meier method and the log-rank test. We assessed group differences using unpaired two-tailed Student’s t-tests (for two groups) or ANOVA (for three or more groups). The results are presented as mean ± SD, and statistical significance was defined as a p-value of less than 0.05.

## Results

### ELAPOR1 is downregulated in the basal-like/squamous PDAC and associates with advanced stage and poor patient survival

We aimed to identify candidate tumor suppressor genes involved in the development of the basal-like/squamous subtype of PDAC. Initially, we analyzed transcriptome data from two PDAC cohorts (Bailey cohort and Moffitt cohort)^3, 4^, which identified 399 genes that were differentially expressed in the basal-like/squamous vs classical/progenitor subtypes of the disease (cutoffs: fold change > 1.5 or < -1.5, p < 0.05), with 268 of these genes being downregulated in the basal-like/squamous subtype (Figure 1A). We further analyzed the genes based on their association with patient survival (hazard ratio < 0.666 using COX regression analysis, p < 0.05) in our NCI-UMD-German patient cohort, yielding a narrowed list of 60 genes (Table S2).

**Figure 1.**
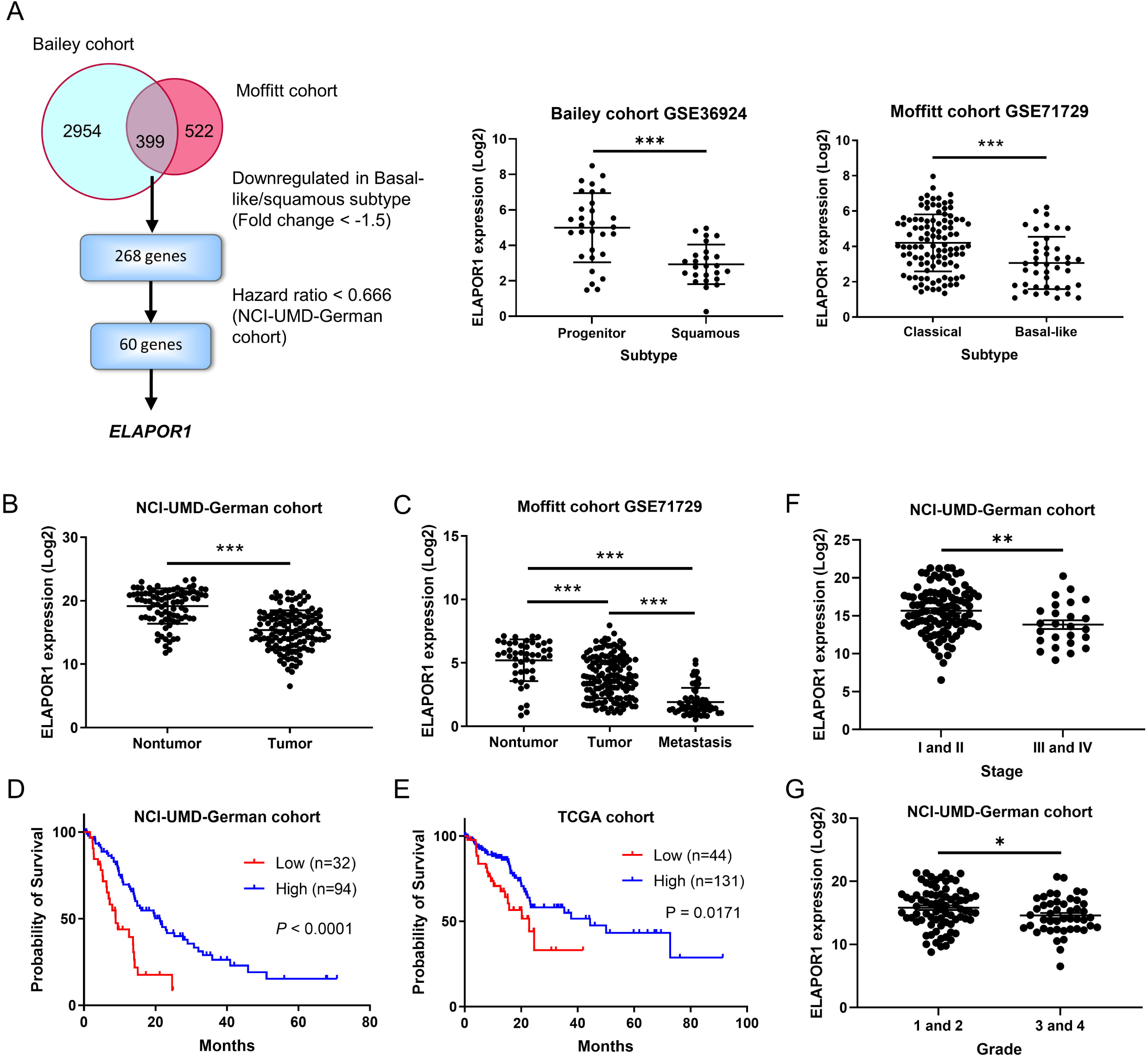
Transcriptome analysis identifies ELAPOR1 as a candidate tumor suppressor in basal-like/squamous PDAC tumors and as a prognostic marker of patient survival. (A) Approach to identify potential suppressor genes of the basal-like/squamous subtype. Candidate genes are identified by analyzing two cohorts (Bailey cohort and Moffitt cohort)^3, 4^, followed by a survival analysis in the NCI-UMD-German cohort. (B-E) *ELAPOR1* transcript levels in PDAC tumors compared to adjacent non-cancerous tissues. *ELAPOR1* is found to be downregulated in tumors in both the NCI-UMD-German cohort and a validation cohort (Moffitt cohort; GSE71729). The lower panels show Kaplan-Meier plots depicting the association between decreased *ELAPOR1* and decreased PDAC patient survival in the NCI-UMD-German and a validation cohort (TCGA cohort). (F and G) Downregulation of *ELAPOR1* is observed in advanced stage (III/IV) and higher grade of PDAC. Data represent mean ± SD. Significance testing with unpaired t-test. *p < 0.05, **p < 0.01, ***p < 0.005

Among them, *IAPP* and *ELAPOR1* were respectively ranked as the first and the second most downregulated gene in tumor tissues compared to nontumor tissues, and *ELAPOR1* was more significantly downregulated in the basal-like/squamous subtype in both the Bailey and Moffitt cohort datasets (Table S2). *ELAPOR1* is an estrogen-regulated gene and has been identified as a tumor suppressor gene in other cancers^24–33^. We hypothesized that ELAPOR1 inhibits PDAC progression through metabolic reprogramming and selected it for further investigation.

Downregulation of *ELAPOR1* transcript levels associated with decreased patient survival, and *ELAPOR1* transcripts were downregulated in tumor tissues compared with nontumor tissues in both the NCI-UMD-German cohort and the validation cohorts (Moffitt cohort and TCGA cohort) (Figure 1B-E). Moreover, the downregulation of *ELAPOR1* associated with advanced stage (III/IV) and higher grade of PDAC (Figure 1F and 1G). ELAPOR1 protein was commonly detected in the cytoplasm of acinar cells and endocrine cells, but less so in tumor cells (Figure S1A and S1B). ELAPOR1 protein found to be downregulated in epithelial cells with acinar-to-ductal metaplasia (ADM) features, suggesting that ELAPOR1 might be involved in maintaining acinar cell differentiation. These findings together support the hypothesis that ELAPOR1 may have a potential tumor inhibitory function and its reduction is associated with poor patient survival and the basal-like/squamous subtype.

### Upregulation of ELAPOR1 downregulates cellular movement in PDAC

We next investigated whether ELAPOR1 expression may decrease basal-like/squamous subtype characteristics and reduce disease aggressiveness. We previously defined the molecular subtypes of 175 patients in our NCI-UMD-German cohort^17^ and showed that pathways related to cellular movement are more activated in the basal-like/squamous than the classical/progenitor subtype.

Tumors defined as basal-like/squamous subtype tended to have decreased *ELAPOR1* mRNA expression when compared to other tumors (Figures S2A). Pathway enrichment analysis using the Ingenuity Pathways Analysis (IPA) revealed that ELAPOR1-high PDAC tumors enriched the pathways, including cellular movement and cellular development (Figure 2A).

**Figure 2.**
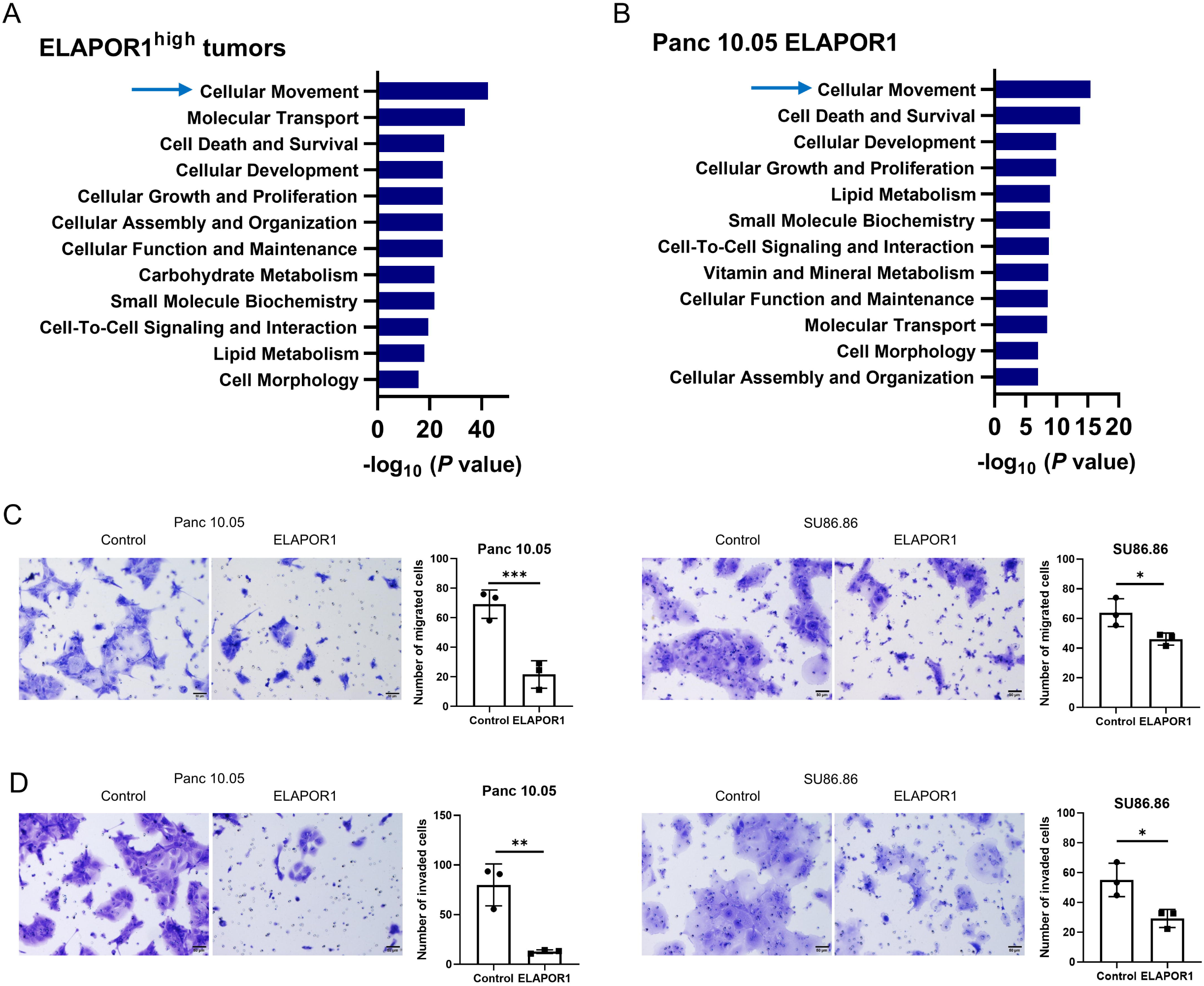
ELAPOR1 upregulation suppresses migration and invasion in PDAC. (A-B) Transcriptomic analyses of patient PDAC and Panc 10.05 human PDAC cells. Ingenuity pathway analysis (IPA) of contrasts *ELAPOR1*-high versus *ELAPOR1*-low PDAC tumors and Pan 10.05 cells with *ELAPOR1* transgene expression (*ELAPOR1*-high) versus vector control (*ELAPOR1*-low). Shown are IPA enrichment scores (*P* value-based) for pathways that are inhibited by increased ELAPOR1. IPA highlights the association of elevated ELAPOR1 with similar pathways in tumors and cultured cells, including down-regulation of cellular movement and cellular development, in *ELAPOR1*-high PDAC tumors and in human PDAC cell lines with *ELAPOR1* transgene overexpression. (C-D) PDAC cells with ELAPOR1 transgene overexpression exhibit decreased migration and invasion ability when compared to the vector control cells. Data represent mean ± SD of 3 replicates with unpaired t-test. *p < 0.05, **p < 0.01, ***p < 0.005.

To confirm these observations, we evaluated ELAPOR1 expression in human PDAC cell lines. ELAPOR1 expression was low to undetectable in all the PDAC cell lines (Figure S2B). Panc 10.05 and SU86.86 cells were then selected to establish cell lines with *ELAPOR1* transgene expression. ELAPOR1 expression was confirmed at the mRNA and the protein levels in these cell lines (Figure S2C and S2D). Pathway enrichment analysis using IPA showed that the human PDAC cells with ELAPOR1 transgene expression had an enrichment pattern in pathways coherent with ELAPOR1-high PDAC tumors of the NCI-UMD-German cohort, showing again an enrichment of genes in the cellular movement pathway (Figure 2B). To further investigate ELAPOR1 functions, we performed additional *in vitro* experiments using these ELAPOR1-overexpressing PDAC cells. CCK-8/WST-8 assay indicated that ELAPOR1 did not affect proliferation (data not shown). However, ELAPOR1 expression significantly inhibited the migration and invasion of PDAC cells (Figure 2C and 2D). Furthermore, IPA and gene set enrichment analysis (GSEA) in PDAC tumors revealed that ELAPOR1 may upregulate neuroendocrine differentiation and downregulate HIF1A/hypoxia signaling (Figure 3A-3B and Figure S3A), which are known characteristics of the classical/progenitor subtype^3^.These findings suggest that ELAPOR1 plays a role in suppressing the characteristics of the basal-like/squamous subtype of PDAC.

**Figure 3.**
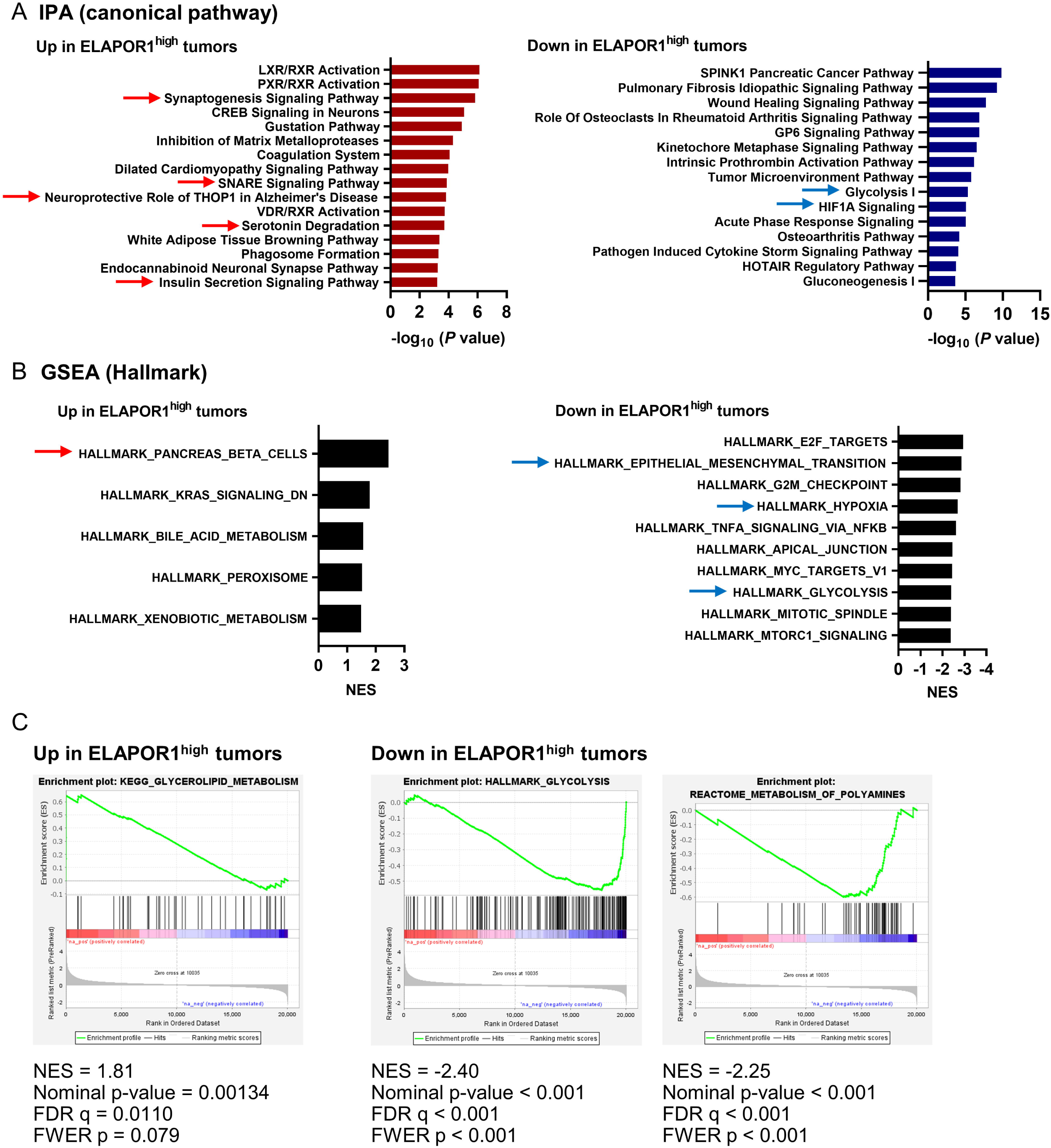
ELAPOR1 upregulation promotes classical/progenitor subtype characteristics and metabolic reprogramming in PDAC. (A-B) IPA and Gene Set Enrichment Analysis (GSEA) using transcriptome data for the contrast *ELAPOR1*-high versus *ELAPOR1*-low PDAC tumors. Combined, the IPA of canonical pathways and GSEA results suggest that high ELAPOR1 activates neuron differentiation pathways and downregulates pathways associated with glycolysis, HIF1A/hypoxia, and epithelial-mesenchymal transition. (C) The GSEA highlights the activation of lipogenesis and the downregulation of glycolysis and polyamine metabolism in ELAPOR1-high PDAC.

### Upregulation of lipid metabolism and downregulation of amino acid metabolism in ELAPOR1-high PDAC

We next investigated the effect of ELAPOR1 on metabolism in PDAC using transcriptome and metabolome analysis from a total of 50 patients in our NCI-UMD-German cohort. Applying GSEA to the transcriptome data revealed that ELAPOR1 expression upregulates lipogenesis while downregulating glycolysis and polyamine metabolism, a downstream pathway of amino acid metabolism (Figure 3C and Figure S3B).

Expanding our analyses by metabolomic analysis, we identified 70 significantly upregulated and 64 downregulated metabolites in *ELAPOR1*-high tumors compared to *ELAPOR1*-low tumors (Figure 4A and Table S3). *ELAPOR1* expression levels in the unclassified and the classical/progenitor subtypes were rather similar (Figure S2A). Thus, we hypothesized that ELAPOR1 is not involved in maintaining differences between these two subtypes and compared metabolic profiles of the unclassified and classical/progenitor subtypes as one dataset with the metabolic profile of the basal-like/squamous subtype. 98 metabolites were significantly increased and 155 were decreased in the combined classical/progenitor and unclassified subtype group when compared to the basal-like/squamous subtype group (Figure S4A and Table S4). Pathway enrichment analysis using MetaboAnalyst 5.0 (https://www.metaboanalyst.ca) with these differential metabolites as input pointed to the upregulation of lipid metabolism and the downregulation of amino acid and carbohydrate metabolism in both ELAPOR1-high tumors and the non-basal-like/squamous tumors (Figure 4B and 4C; Figure S4B and S4C). Furthermore, the enrichment analysis using IPA with the upregulated metabolites in both the *ELAPOR1*-high tumors and the non-basal-like/squamous tumors indicated the activation of neuronal pathways (Figure S4D and S4E).

**Figure 4.**
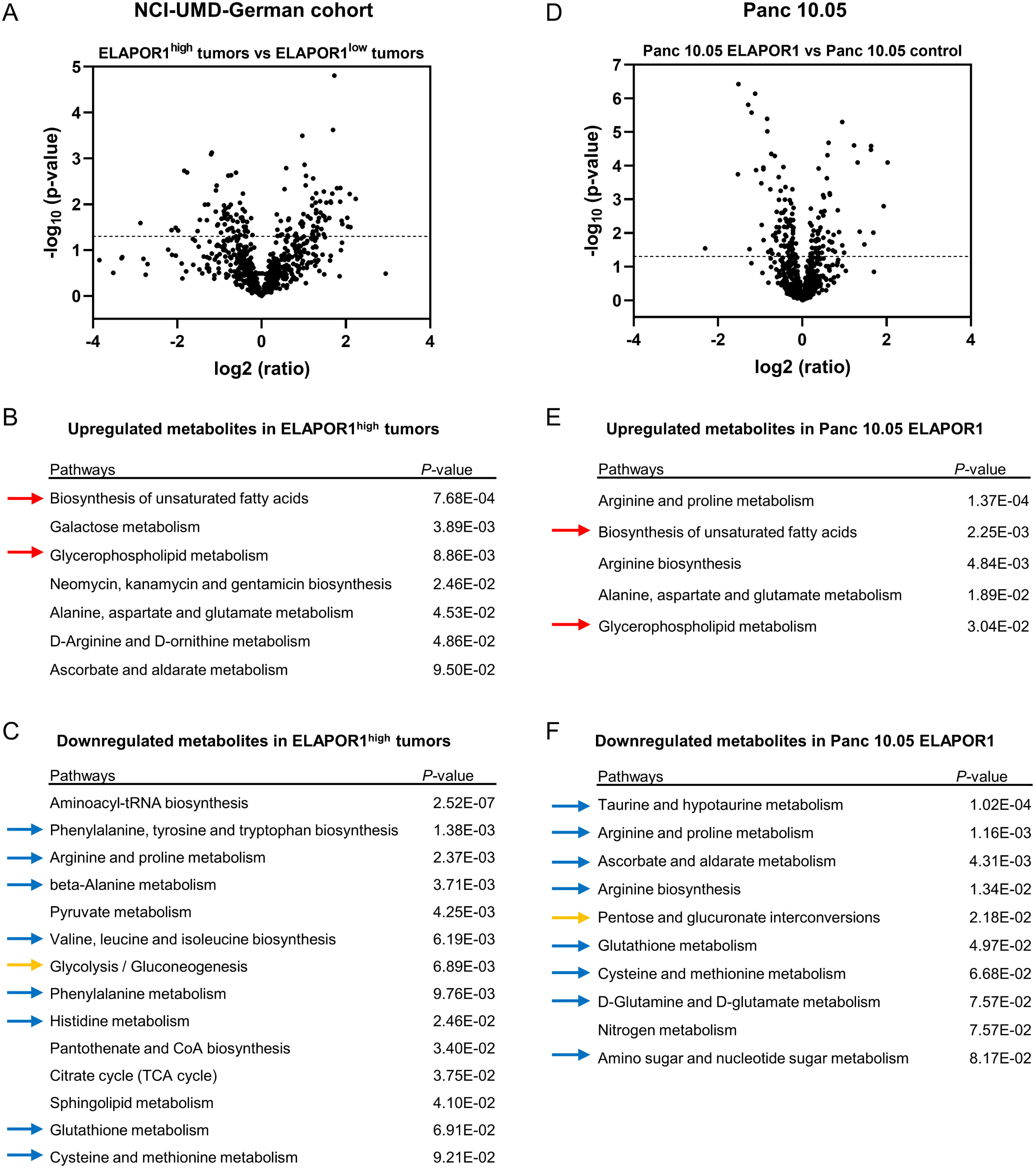
Upregulation of lipid metabolism and downregulation of amino acid metabolism in ELAPOR1-high PDAC. Analysis of the metabolome in PDAC tumors (NCI-UMD-German cohort) and Pan 10.05 human PDAC cells, contrasting *ELAPOR1*-high (n = 25) versus *ELAPOR1*-low (n = 25) PDAC tumors and Pan 10.05 cells with *ELAPOR1* transgene expression (*ELAPOR1*-high) versus vector control (*ELAPOR1*-low). (A) Volcano plot of the metabolites with differential abundance in ELAPOR1-high versus ELAPOR1-low tumors. 70 metabolites are significantly upregulated and 64 are downregulated in ELAPOR1-high tumors (p < 0.05). The dotted line indicates -log10 (p = 0.05). (B-C) Pathway enrichment analysis with differential metabolites using MetaboAnalyst 5.0, indicating that lipogenesis-related pathways are upregulated (red arrows) and amino acid (blue arrows) and carbohydrate metabolism (orange arrow) are down-regulated in ELAPOR1-high tumors. (D) Volcano plot of the metabolites with differential abundance in ELAPOR1-overexpressing versus vector control Panc 10.05 cells. 64 metabolites are significantly upregulated and 85 are downregulated in ELAPOR1-overexpressing Panc 10.05 cells. The dotted line indicates -log10 (p = 0.05). (E and F) Pathway enrichment analysis with differential metabolites using MetaboAnalyst 5.0, indicating that ELAPOR1-overexpressing Panc 10.05 cells exhibit upregulated lipogenesis (red arrows) and downregulated amino acid (blue arrows) and carbohydrate metabolism (orange arrow).

To continue our investigation of the role of these metabolites, we performed a global metabolome analysis in PDAC cells. Here, we observed that 64 metabolites were significantly upregulated and 85 were downregulated in *ELAPOR1* transgene-overexpressing Panc 10.05 cells when compared to vector control cells (Figure 4D and Table S5). Pathway enrichment analysis using MetaboAnalyst 5.0 revealed that ELAPOR1-overexpressing Panc 10.05 cells upregulated lipogenesis, while downregulating amino acid and carbohydrate metabolism (Figure 4E and 4F). Previous studies from our group have shown that amino acid metabolism promotes the aggressiveness of PDAC^17^, while lipogenesis is inhibiting disease progression^34^. We observed similar metabolic adaptation profiles in basal-like/squamous PDAC and ELAPOR1-low PDAC cells, which may potentially contribute to the disease progression.

## 1-Methylnicotinamide accelerates migration and invasion in PDAC cells

Twelve metabolites were significantly upregulated and 15 were downregulated in both ELAPOR1-high PDAC tumors and ELAPOR1-overexpressing Panc 10.05 cells (Table S6). N-acetyl-aspartyl-glutamate (NAAG), N-acetylaspartate (NAA), guanidinoacetate, docosahexaenoate (DHA), and ornithine were highly upregulated in ELAPOR1-overexpressing Panc 10.05 cells (p < 0.05, ratio > 1.5, Figure 5A). Notably, NAAG and NAA are metabolites with neurotransmitter function, suggesting that ELAPOR1 promotes neuronal differentiation in cellular metabolism. Conversely, 1-methylnicotinamide (MNA), myristoyl dihydrosphingomyelin, N-palmitoyl-sphinganine, palmitoyl dihydrosphingomyelin, stearoylcarnitine, and 3-hydroxylaurate were highly downregulated in ELAPOR1-overexpressing Panc 10.05 cells (p < 0.05, ratio < 0.666, Figure 5B), with MNA showing the most striking downregulation. These metabolites were also downregulated in the combined classical/progenitor and unclassified PDAC group (Figure S5). MNA is a metabolite originating from an amino acid, S-adenosylmethionine. In the NCI-UMD-German cohort, we found a strong correlation between MNA and methionine levels in the PDAC tumors (p < 0.0001, r = 0.7411, Figure S4F). S-adenosylmethionine originates from methionine. Previous study has shown that MNA inhibits interferon gamma in ovarian cancer, leading to tumor progression^35^. In our study, GSEA showed enhanced interferon gamma signaling in MNA-low ELAPOR1-overexpressing Panc 10.05 cells (Figure S3C). Hence, we hypothesized that the downregulation of amino acid metabolism in ELAPOR1-high tumors may result in a decrease of MNA, leading to decreased disease aggressiveness in PDAC patients. Consistent with the hypothesis, exposure of PDAC cells to MNA resulted in enhanced migration and invasion (Figure 5C and 5D). These findings suggest that ELAPOR1-induced metabolic alterations may have a potential role in reducing disease aggressiveness particularly in classical subtype.

**Figure 5.**
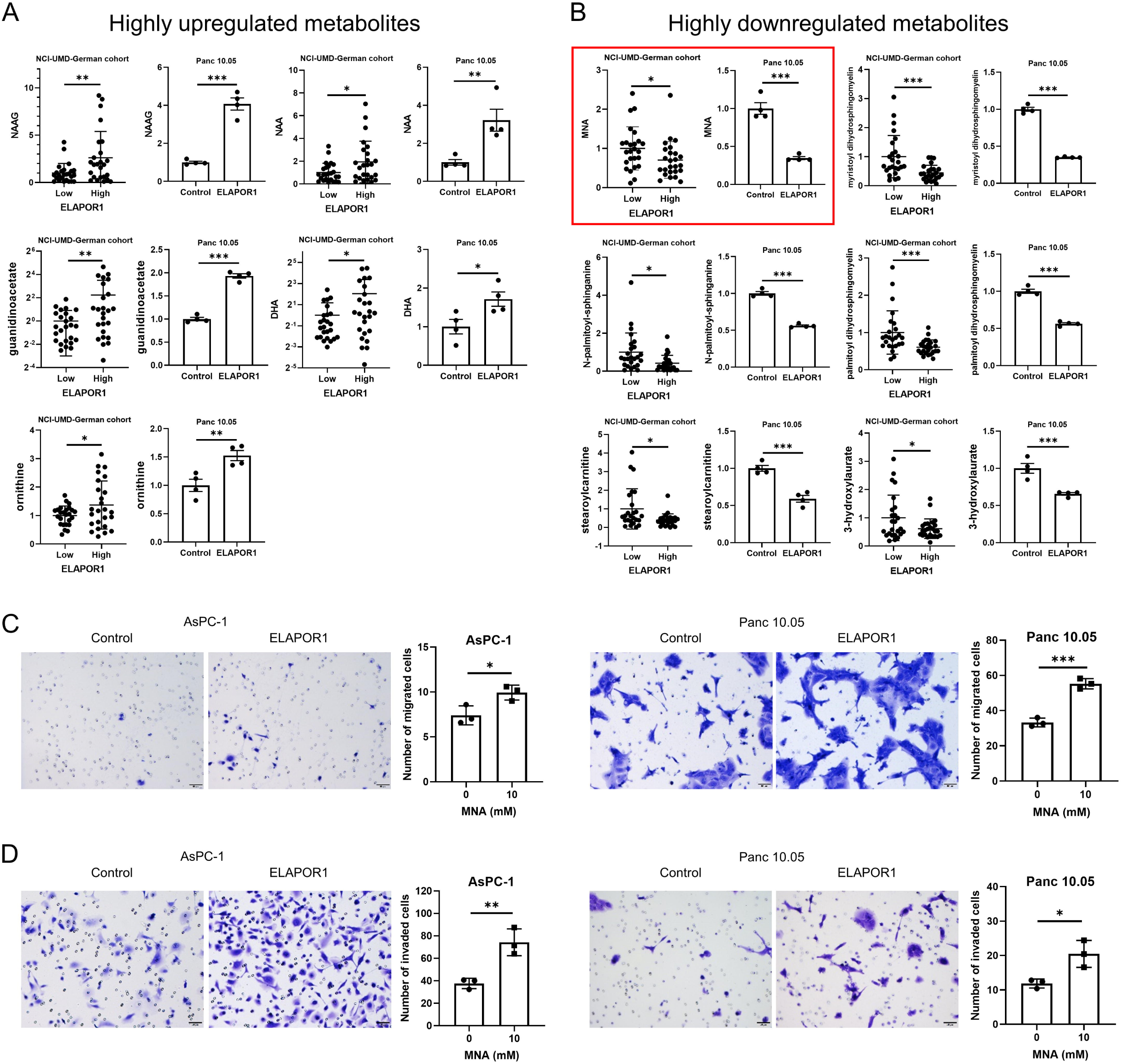
1-Methylnicotinamide promotes migration and invasion in PDAC cells. (A) Examples of metabolites highly upregulated in both ELAPOR1-high PDAC tumors (NCI-UMD-German cohort) and ELAPOR1 transgene expressing Panc 10.05 cells. (B) Metabolites highly downregulated in both ELAPOR1-high PDAC tumors and ELAPOR1 transgene expressing Panc 10.05 cells. (C-D) MNA promotes migration and invasion in cultured human PDAC cells. Data represent mean ± SD with unpaired t-test. *p < 0.05, **p < 0.01, ***p < 0.005. MNA; 1-methylnicotinamide, NAAG; N-acetylaspartylglutamate, NAA; N-acetylaspartate, DHA; docosahexaenoic acid.

## Discussion

In this study, we observed that downregulation of ELAPOR1 associates with the basal-like/squamous subtype and an unfavorable patient prognosis in PDAC. Upregulation of ELAPOR1 induced changes to cancer cell metabolism and inhibited migration and invasion in PDAC cells. Moreover, ELAPOR1 expression associated with neuroendocrine differentiation, a characteristic of the classical/progenitor subtype in which ELAPOR1 expression is maintained.

*ELAPOR1*, also known as *KIAA1324* or estrogen-induced gene 121 (*EIG121*), encodes a transmembrane protein that localizes to various cellular membranes, including the plasma membrane, lysosome, endosome, and Golgi apparatus^24, 25^. ELAPOR1 regulates secretory granule differentiation^25^ and autophagy under stress conditions^24, 36, 37^. In gastric cancer, ELAPOR1 suppresses disease progression^32^. Autophagy plays a critical role in cellular metabolism and has both promoting and suppressing effects in cancer^38^, and may suppress the initiation of PDAC^38–40^. In pancreatic β cells, ELAPOR1 inhibits insulin hormone and insulin-like growth factor 1 signaling by decreasing the number of cell-surface receptors and blunting the response to insulin^26^. In tumors, the expression of ELAPOR1 has been shown to be a favorable prognostic biomarker in prostate, ovarian, endometrial, pancreatic neuroendocrine, and breast cancer^27–31^. ELAPOR1 also suppresses the progression of gastric cancer by inhibiting the oncoprotein GRP78 ^32^. However, the function of ELAPOR1 in breast cancer remains controversial^24, 36, 37^. A link between ELAPOR1 and metabolism has been reported^33^; however, the role of ELAPOR1 in PDAC have not been described.

Cancer cells increase glycolysis, and lipid and amino acid synthesis through the induction of the pentose phosphate pathway^8, 9^. Previous studies showed that the basal-like/squamous subtype relies upon upregulated glycolysis and amino acid metabolism, while the classical/progenitor subtype associates with increased lipogenesis^13–17^. Amino acids are vital metabolites for PDAC progression^13, 41, 42^, while lipids may have more complex effects^34, 42, 43^.

For example, we previously reported that fatty acids exert growth inhibitory effects in PDAC^34^. Consistently, ELAPOR1 induced alterations to the metabolism may promote the classical/progenitor subtype because of the downregulation of carbohydrate and amino acid metabolism and the upregulation of lipogenesis. However, its loss may redirect differentiation into the basal-like/squamous subtype with an increase in glycolysis and decrease in lipogenesis.

As a metabolite of interest, MNA was one of the most strikingly down-regulated metabolites in ELAPOR1-overexpressing Panc 10.05 cells. MNA has been described as an immune regulatory metabolite in human ovarian cancer^35^. MNA inhibits interferon gamma signaling in ovarian cancer, leading to tumor progression^35^. We found enhanced interferon gamma signaling in MNA-low ELAPOR1-overexpressing Panc 10.05 cells, suggesting similar functions of MNA in ovarian cancer and PDAC. Other metabolites, like NAA, NAAG, guanidinoacetate, DHA, and ornithine were also highly upregulated in ELAPOR1-overexpressing Panc 10.05 cells. Moreover, the function of most of these metabolites has not been described in PDAC. NAA and NAAG are metabolites with neurotransmitter function, suggesting that ELAPOR1 promotes neuronal differentiation through cellular metabolism. These and other observations should be followed up in future studies.

Molecular subtyping in breast cancer has undergone extensive research and has proven to be clinically valuable^44^. Breast cancer is commonly classified into four distinct molecular subtypes. Diagnosis of the molecular subtype in each patient’s breast cancer is typically determined using IHC, enabling the selection of appropriate therapeutic agents based on the diagnosis. Conversely, molecular subtyping in PDAC is still in its exploratory phase and PDAC molecular subtypes are currently being investigated using whole exome sequencing methods such as RNA sequencing. However, this study has made an important observation, identifying ELAPOR1 as a potential diagnostic marker for the molecular subtype of PDAC. If the molecular subtype of PDAC can be readily diagnosed using techniques such as qPCR or IHC, it may have significant clinical implications.

Taken together, our data suggest that persistence of ELAPOR1 expression promotes transcriptomic and metabolomic characteristics indicative of the classical/progenitor subtype, whereas its loss associates with basal-like/squamous tumors and increased aggressiveness of PDAC. ELAPOR1 expression induced a distinct metabolic signature characterized by upregulation of lipogenesis and downregulation of amino acid metabolism, commonly observed in the classical/progenitor PDAC subtype. Upregulation of ELAPOR1 also associated with improved PDAC survival and exerted inhibitory effects on migration and invasion in PDAC cells, indicating its potential tumor inhibitory function and may be further explored for designing novel interventional approaches in PDAC patients.

## Author Contributions

**Yuuki Ohara:** Conceptualization, Methodology, Resources, Investigation, Writing – Original Draft, Writing – Review & Editing, Project Administration. **Amanda J. Craig**: Methodology, Data Curation, Writing – Review & Editing, Project Administration. **Huaitian Liu:** Data Curation, Writing – Review & Editing, Project Administration. **Shouhui Yang:** Methodology, Writing – Review & Editing, Project Administration. **Paloma Moreno:** Project Administration. **Tiffany H. Dorsey:** Project Administration. **Helen Cawley:** Project Administration. **Azadeh Azizian:** Resources, Project Administration. **Jochen Gaedcke:** Resources, Project Administration. **Michael Ghadimi:** Resources, Project Administration. **Nader Hanna:** Resources, Project Administration. **Stefan Ambs:** Methodology, Writing – Review & Editing. **S. Perwez Hussain:** Conceptualization, Writing – Review & Editing, Supervision, Funding Acquisition. The work reported in the paper has been performed by the authors, unless clearly specified in the text.

### Abbreviations

(MNA): 1-methylnicotinamide
(ADM): acinar-ductal metaplasia
(ADEX): aberrantly differentiated endocrine exocrine
(DHA): docosahexaenoate
(ELAPOR1): endosome-lysosome associated apoptosis and autophagy regulator 1
(EIG121): estrogen induced gene 121
(GSEA): Gene Set Enrichment Analysis
(HIF): hypoxia-inducible factor
(IHC): immunohistochemistry
(IPA): Ingenuity pathway analysis
(NAA): N-acetylaspartate
(NAAG): N-acetyl-aspartyl-glutamate
(PDAC): pancreatic ductal adenocarcinoma
(STR): short tandem repeat

## Acknowledgments

Authors would like to thank staff and study coordinators at the University of Maryland School of Medicine who were involved in the procurement of clinical biospecimens and patient data. We would also like to thank the staff of the Department of General, Visceral and Pediatric Surgery, University Medical Center Göttingen, Göttingen, Germany for their help with clinical samples. Lastly, we would like to thank the staff of the NCI-CCR Sequencing Facility Frederick for their contributions.

## Data and materials availability

Data and materials are available upon request.

The RNA sequencing data were deposited in the NCBI’s Gene Expression Omnibus (GEO) database under accession number (GSE243879). GSE243879: ELAPOR1 induces the classical/progenitor subtype and contributes to reduced disease aggressiveness through metabolic reprogramming in pancreatic cancer

To protect the confidentiality of our data and results, the repository will remain private while the manuscript is under review.

## Ethics Statement

Pancreatic tissues were collected from resected PDAC patients at the University of Maryland Medical System (UMMS) in Baltimore, MD, through an NCI-UMD resource contract, and at the University Medical Center Göttingen, Germany. Board-certified pathologists evaluated PDAC histopathology. The use of these clinical samples for our study was reviewed by the NCI-Office of the Human Subject Research Protection (OHSRP) at the NIH in Bethesda, MD (Exempt#4678). All participants provided written informed consent. The conducted procedures adhered to ethical standards and were in accordance with the 1975 Declaration of Helsinki, as revised in 2008.

## Supplementary figure legends

**Figure S1.**
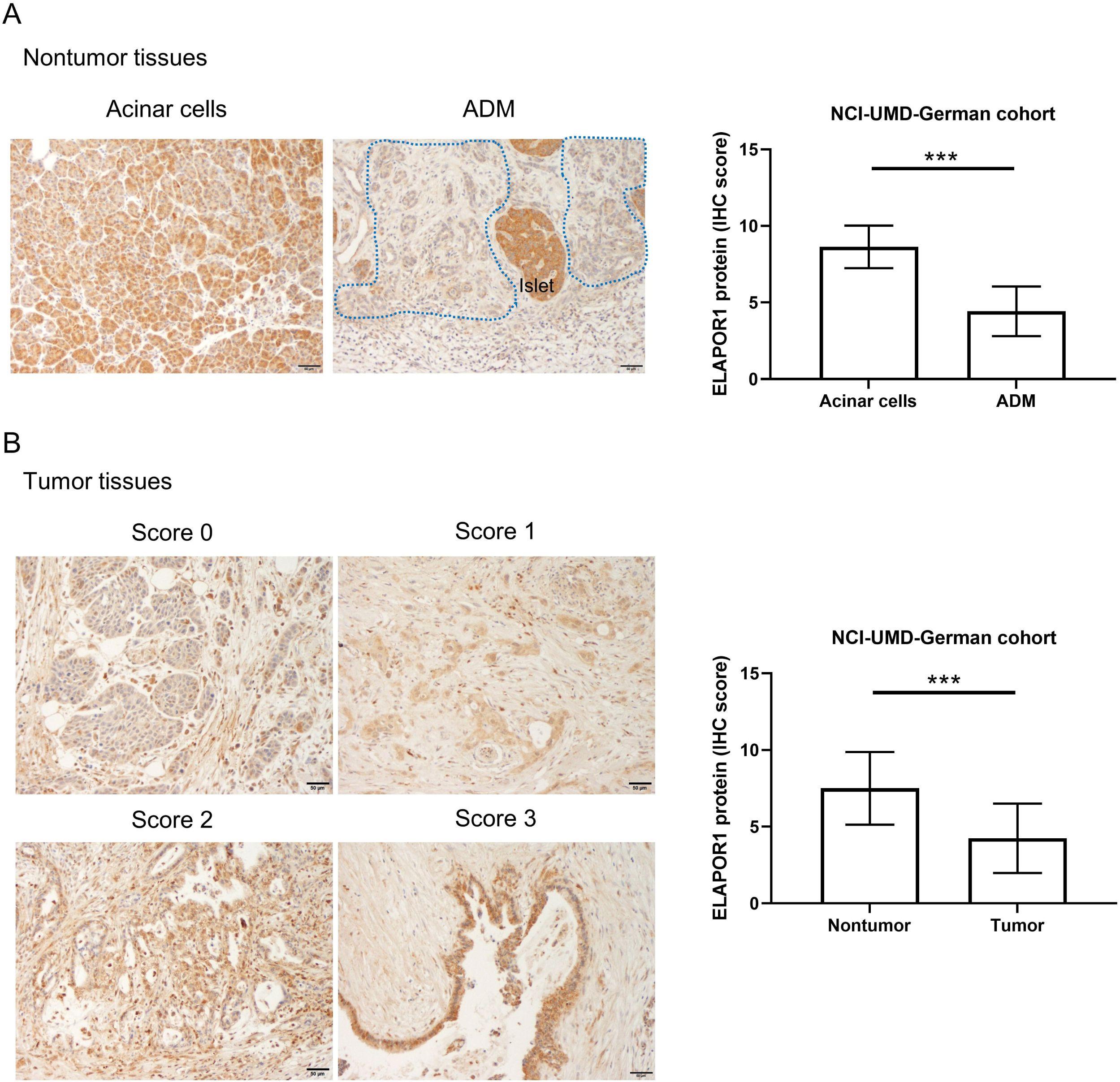
Downregulation of ELAPOR1 protein in PDAC tumors in the NCI-UMD-German cohort. IHC of ELAPOR1 in tumor and non-tumor tissue sections of PDAC patients. (A) ELAPOR1 protein is detected in the cytoplasm, as shown by the brown DAB-based IHC in the tumor cells, the acinar cells, and the endocrine cells in pancreatic islets. Representative ELAPOR1 protein staining in non-cancerous acinar cells (intensity score 3) and acinar-to-ductal metaplasia (ADM, within dotted lines) (intensity score 1), highlighting a significant downregulation of ELAPOR1 in ADM. (B) ELAPOR1 in representative tumor sections (scores 0-3), indicating global downregulation of ELAPOR1 in tumor cells when compared to the nontumor acinar cells (see graph to the right). The staining strength is categorized as follows: score 0 (non-stained), score 1 (weak), score 2 (moderate), and score 3 (strong). Scale bar is 20 μm. More details can be found in Materials and Methods. Data represent mean ± SD. Significance testing with unpaired t-test. *p < 0.05, **p < 0.01, ***p < 0.005. ADM; acinar-to-ductal metaplasia, DAB; 3, 3’-diaminobenzidine, IHC; immunohistochemistry.

**Figure S2.**
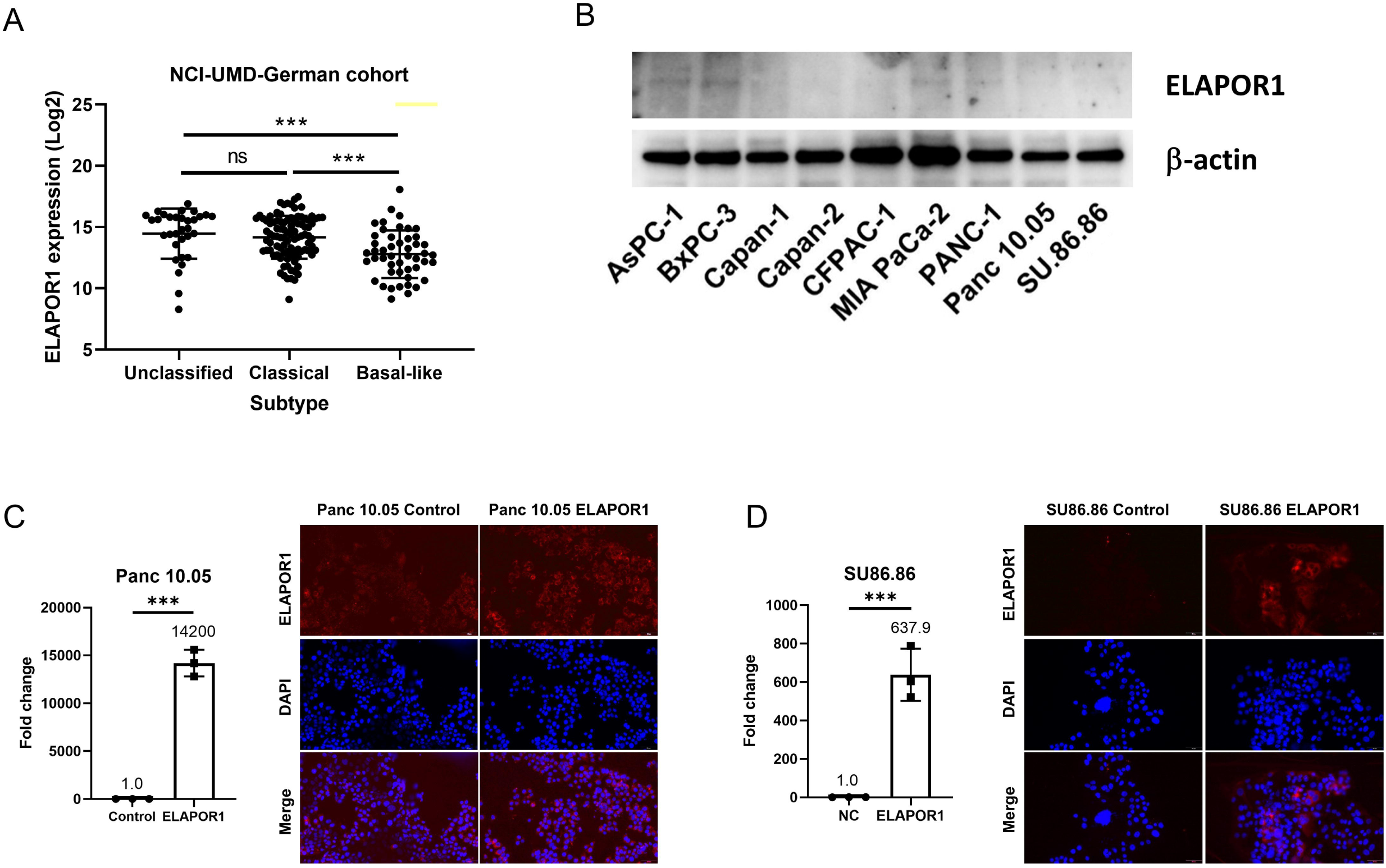
ELAPOR1 expression in human PDAC. (A) ELAPOR1 mRNA expression by subtype in the NCI-UMD-German cohort. (B) Endogenous levels of ELAPOR1 protein in various human PDAC cell lines, revealing consistent suppression of ELAPOR1 expression across all examined PDAC cell lines. (C-D) Confirmation of ELAPOR1 transgene overexpression at the mRNA (bar graph) and protein levels in Panc 10.05 and SU86.86 cells. Data represent mean ± SD of 3 replicates with unpaired t-test. *p < 0.05, **p < 0.01, ***p < 0.005.

**Figure S3.**
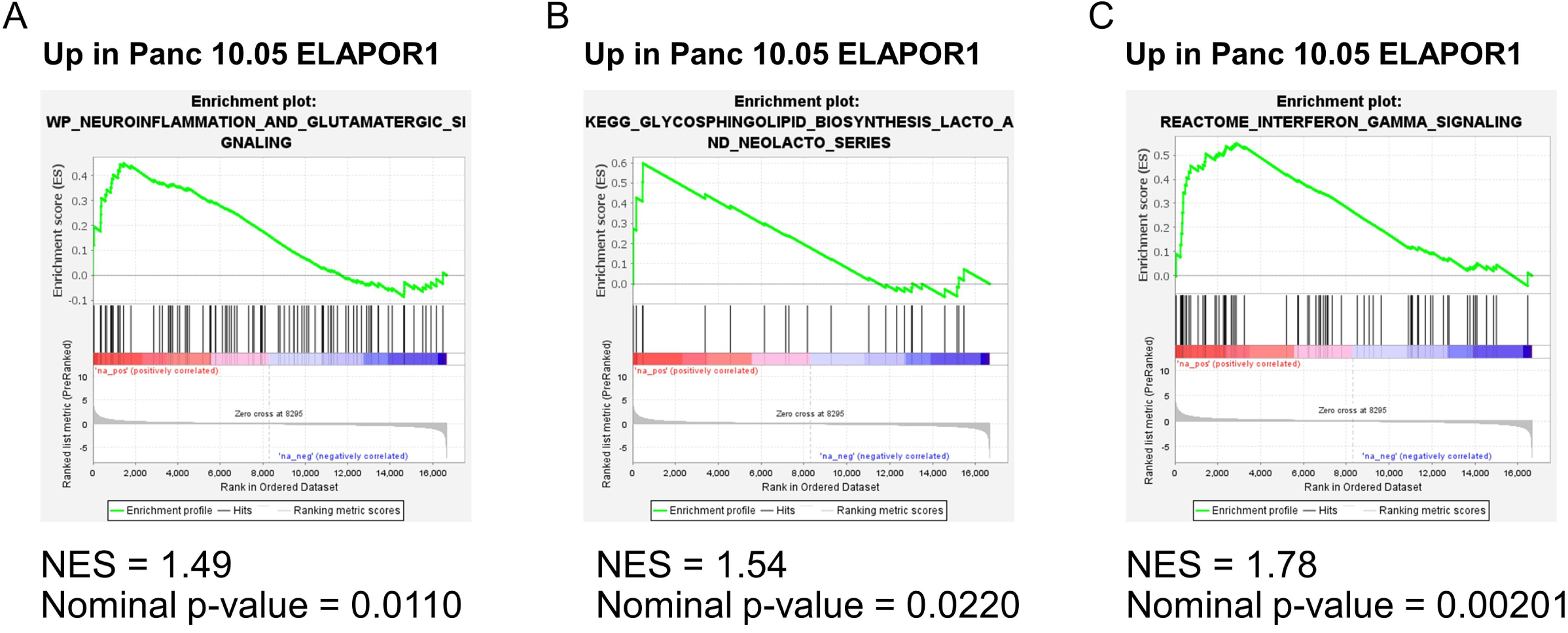
Upregulation of ELAPOR1 promotes neuron differentiation and metabolic reprogramming in PDAC cells. Transcriptomic analysis in Panc 10.05 cells with ELAPOR1 transgene overexpression. The GSEA highlights the activation of neuronal signaling (A), lipogenesis (B), and interferon-gamma signaling (C).

**Figure S4.**
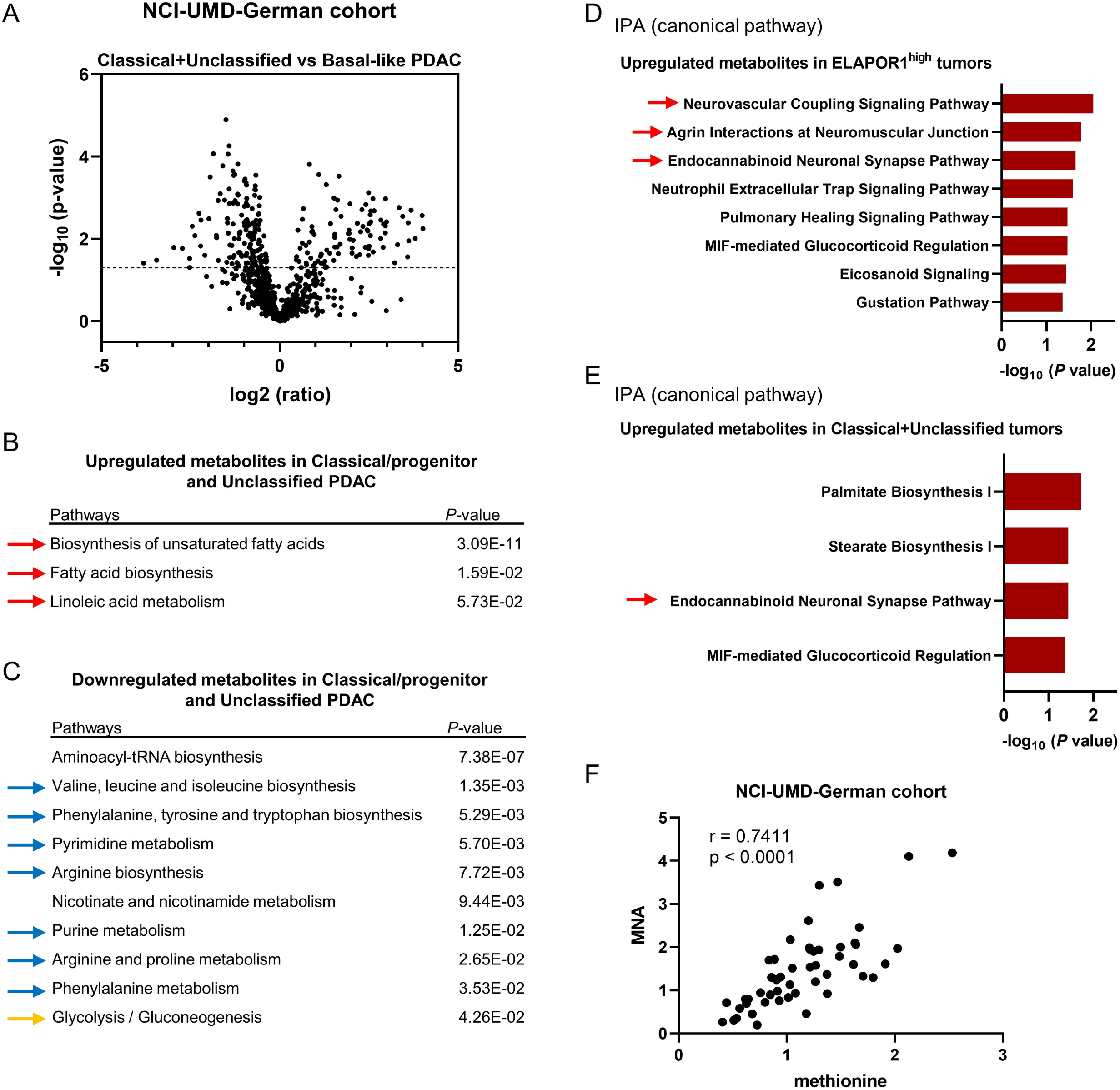
Upregulation of lipid metabolism and downregulation of amino acid metabolism in the classical/progenitor and unclassified PDAC. Metabolome analyses of tumors from PDAC patients in the NCI-UMD-German cohort, contrasting classical/progenitor and unclassified tumors (n = 35) versus basal-like (n = 15) tumors. (A) Volcano plot of the metabolites with differential abundance comparing classical + unclassified versus basal-like/squamous PDAC. 98 metabolites are significantly upregulated and 155 are downregulated in the classical + unclassified PDAC tumors when compared with basal-like tumors. The dotted line indicates -log10 (p = 0.05) thresholds. (B-C) Pathway enrichment analysis using MetaboAnalyst 5.0, indicating that the classical/progenitor + unclassified PDAC subtype group shows upregulation of lipogenesis (red arrows) and downregulation of amino acid (blue arrows) and carbohydrate metabolism (orange arrow). (D-E) Pathway analysis using IPA, indicating that ELAPOR1-high PDAC and the classical/progenitor + unclassified PDAC may similarly activate neuronal signaling pathways. (F) Pearson’s correlation coefficient, revealing a strong correlation between the abundance levels of MNA and methionine in PDAC tumors. IPA; Ingenuity pathway analysis, MNA; 1-methylnicotinamide.

**Figure S5.**
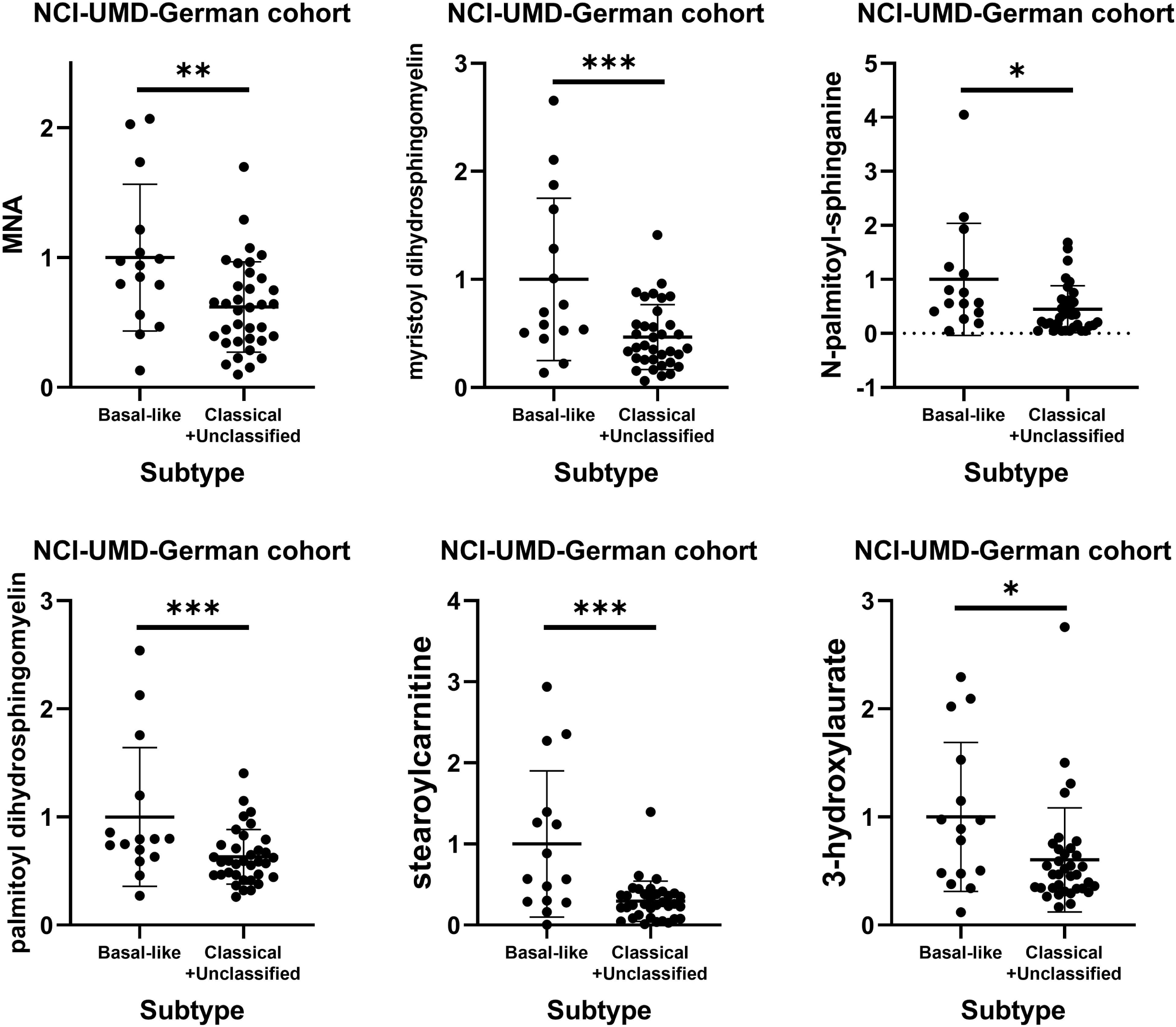
Highly downregulated metabolites in the classical/progenitor and unclassified PDAC. Highly downregulated metabolites in ELAPOR1-high patient PDAC tumors and in ELAPOR1-overexpressing Panc 10.05 cells (Figure 5) are also downregulated in the classical/progenitor + unclassified PDAC (classical/progenitor + unclassified PDAC; n = 35, basal-like/squamous PDAC; n = 15).

